# Cryopreservation and transplantation of common carp spermatogonia

**DOI:** 10.1101/429449

**Authors:** Roman Franěk, Zoran Marinović, Jelena Lujić, Béla Urbányi, Michaela Fučíková, Vojtěch Kašpar, Martin Pšenička, Ákos Horváth

## Abstract

Common carp (*Cyprinus carpio*) is one of the most cultured fish species over the world with many different breeds and plenty of published protocols for sperm cryopreservation, however, data regarding preservation of gonadal tissue and surrogate production is still missing. A protocol for freezing common carp spermatogonia was developed through varying different factors along a set of serial subsequent experiments. Among the six cryoprotectants tested, the best survival was achieved with dimethyl sulfoxide (Me_2_SO). In the next experiment, a wide range of cooling rates (0.5–10 °C/min) and different concentrations of Me_2_SO were tested resulting in the highest survival using 2 M Me_2_SO and cooling rate of –1 Q59 A When testing different tissue sizes and incubation times in the cryomedium, the highest viability was observed when incubating 100 mg tissue fragments for 30 min. Finally, sugar supplementation did not yield significant differences. When testing different equilibration (ES) and vitrification solutions (VS) used for needle-immersed vitrification, no significant differences were observed between the tested groups. Additionally, varied exposure time to VS did not improve the vitrification outcome where the viability was 4-fold lower than that of freezing. The functionality of cryopreserved cells was tested by interspecific transplantation into sterilized goldfish recipients. The exogenous origin of the gonads in goldfish recipients was confirmed by molecular markers and incorporation rate was over 40% in both groups at 3 months post transplantation. Results of this study can serve as an alternative way for long-term preservation of germplasm in carp which can be recovered in a surrogate recipient.

## Introduction

Common carp (*Cyprinus carpio*) is one of the oldest domesticated fish species in the world and is mainly cultured in Europe and Asia. Nowadays, the common carp expanded to all continents with exception of Antarctica. Overall carp production from aquaculture in 2014 was more than 4 million tons – around 14% of the total freshwater aquaculture production in that year [1]. Fruitful history and lasting popularity of this species gave to rise many different strains and lines which became important for breed management and production of hybrids in Central Europe [2–4]. Due to this fact, significant efforts have been committed to preservation of carp genetic resources. Long-term cultivation of pure-breed livestock [5], methods for genetic diversity identification [6–9] as well as methods for creating and keeping gene banks through sperm cryopreservation [10–14] have been developed. However, *ex situ* preservation of valuable genetic material still relies only on sperm cryopreservation.

Recent progress in biotechnology revealed a very efficient alternative for preservation and restoration of a valuable genetic material using cryopreservation of germline stem cells and surrogate offspring production. Germline stem cells (including spermatogonial stem cells –SSCs, oogonial stem cells – OSCs and primordial germ cells – PGCs) have the ability to incorporate into gonads of suitable recipients after transplantation, and therein start gametogenesis and give rise to functional gametes and subsequently offspring. They exhibit a high level of sexual plasticity as transplanted spermatogonia and oogonia can develop into both eggs and sperm, thus both sexes can be easily restored even from a single donor using host individuals of different sex [15–19]. Additionally, several studies so far have displayed that fish germ cells can be cryopreserved, stored for virtually indefinite periods of time, and can be transplanted as they retain their colonization and proliferation capabilities after thawing [20–22].

Most of the cryopreservation studies until now focused on optimizing protocols by freezing germ cells slowly (usually at ∼ 1 °C/min) to a temperature that is optimal for plunging into liquid nitrogen. These studies demonstrated that certain differences and peculiarities can be found in optimal cryopreservation protocols between species, thus, it is necessary to optimize the cryopreservation procedure for each fish species [23–25]. Performed studies focused on optimizing cryoprotectants, their concentrations or additional sugar or protein supplementation, however, little attention was paid to the size of the cryopreserved tissue pieces, incubation time in cryomedium prior freezing and especially the rate of freezing.

Vitrification as an ultra-fast cryopreservation method has been largely overlooked until recently. The studies of Lujić et al. (2017) [26], Seki et al. (2017) [27], Higaki et al. (2017)[28], Marinović et al. (2018) [29] pioneered in developing vitrification methods for fish germ cells as this method has several advantages over the traditional slow-rate freezing: (1) cost efficiency as it does not require special and costly equipment; (2) time-effectiveness as the sample preparation is quicker; (3) low volumes of liquid nitrogen are needed. As high cryoprotectant concentrations are needed to reach the amorphous glassy state of the tissue during vitrification, optimization of the tradeoff between the cryoprotectant combinations, their concentrations and attaining the highest possible cooling rates is of utmost importance.

In the present study we optimized the freezing and vitrification protocols for the common carp spermatogonia and showed possible gaps for improvement which can be generally adopted for cryopreservation of other fish species. Additionally, we demonstrated the functionality of the cryopreserved cells through interspecific spermatogonia transplantation using sterilized goldfish host.

## Material and methods

### Animal ethics

The study was partly conducted at the Department of Aquaculture, Szent István University, Gödöllő,Hungary, and partly at the aquaculture facility of the Genetic Fisheries Center and Laboratory of Germ Cells at the Faculty of Fisheries and Protection of Waters, University of South Bohemia in České Budějovice, Vodňany, Czech Republic. Experiments conducted in Hungary were approved under the Hungarian Animal Welfare Law (Act XXVIII/1998 of the Hungarian Parliament on the protection and humane treatment of animals) by the Government Office of Pest County (approval number: PE/EA/188–6/2016). Experiments conducted in Czech Republic were approved Ministry of Agriculture of the Czech Republic (reference number: 53100/2013-MZE-17214). The methodological protocol of the current study was approved by the expert committee of the Institutional Animal Care and Use Committee (IACUC) of the University of South Bohemia in České Budějovice, Faculty of Fisheries and Protection of Waters (FFPW) in Vodňany according to the law on the protection of animals against cruelty (Act no. 246/1992 Coll., ref. number 16OZ19179/2016–17214). The study did not involve endangered or protected species. Martin Pšenička and Vojtěch Kašpar (CZ01652) own the certificate (CZ 00673) giving capacity to conduct and manage experiments involving animals according to section 15d paragraph 3 of Act no. 246/1992 Coll.

## Cryopreservation experiment

### Testicular tissue preparation

Common carp males (age 1+ year, BW: 128 ± 34 g) used for development and optimization of the cryopreservation protocol were held in a recirculation system at the Department of Aquaculture, Szent István University, Hungary. Fish were kept at a constant temperature of 24 ± 1 °C, fed twice per day with a low-fat diet. Fish were euthanized by an overdose of 2-phenoxyethanol and decapitated, testes were aseptically excised, washed in phosphate buffered saline (PBS) and cleaned of large blood vessels and adjacent connective tissue. Testes were then cut into small fragments, approximately weighing 50 mg, 100 mg or 150 mg (depending on the experiment). Three fragments were used as a fresh control.

### Dissociation procedure

Each tissue fragment was weighed before dissociation, and subsequently transferred into the dissociation medium (L-15 supplemented with 2 mg/ml collagenase, 1.5 mg/ml trypsin and 40 μg/ml of DNase I), minced into small pieces and incubated for 1.5 h at room temperature (RT; 22 °C) on a shaking plate. Digestion was terminated by addition of 10% Fetal bovine serum (FBS) (v/v). In order to obtain a single cell suspension, samples were filtered through 30 µm filters (Sysmex, Germany) and centrifuged for 10 min at 200 ×g. The supernatant was removed and the pellet was resuspended by a gentle pipetting with addition of appropriate volume of L-15 medium.

### Viability assessment

Cell viability was determined by trypan blue differential staining where the dead cells were stained blue while live cells remained unstained. The number of live early stage germ cells was counted in 20 fields of a Bürker-Türk counting chamber for each sample under a light microscope with phase contrast (Nikon Eclipse E600) at 40× magnification. Final cell survival rate was assessed as the percentage of live cells isolated from cryopreserved tissue compared to the number of live cells isolated from the fresh tissue while correcting for the tissue size according to Lujić et al. (2017): *Viability*(%) = (*N_cryopreserved_* / *N_fresh_*) × *CF* × 100 where *CF* = *Weight_fresh tissue_*/*Weight_cryopreserved tissue_*) 26].

### Freezing of testicular fragments

Cryomedia used in the study were composed of cryoprotectants (type and concentration depending on the experiment), 1.5% Bovine serum albumin (BSA), 25 mM Hepes and sugar supplementation (type and concentration depending on the experiment) diluted in PBS. Tissue fragments were loaded into 1.8 ml cryotubes (Nunc^®^) filled with 1 ml of cryomedium. Samples were equilibrated for 15 min or 30 min (depending on the experiment) on ice, and subsequently placed into CoolCell (Biocision) freezer boxes and into a deep freezer (- 80 °C) which enabled cooling rates of ∼ 1 °C/min or were frozen using a Controlled rate freezer (IceCube 14S programmable freezer (IceCube Series v. 2.24, Sy-Lab, Neupurkersdorf, Austria)) set to different cooling rates depending on the test group (see below). After 4 h, samples were plunged into liquid nitrogen for at least 1 day of storage. Samples were thawed in a 26 °C water bath and tissue fragments were washed three times in L-15 where they remained until further work (not longer than 15 min). Digestion and counting procedures were conducted as mentioned above.

Optimization of the freezing protocol was conducted in four sequential experiments where in each experiment one cryopreservation parameter was changed, and the best outcome was used in the subsequent experiment. Initially, 0.1 M glucose was used as sugar supplementation and 100-mg tissue fragments were frozen. Firstly, the effects of dimethyl sulfoxide (Me_2_SO), ethylene glycol (EG), glycerol (Gly), 1:1 combination of Me_2_SO and propylene glycol (Me_2_SO + PG) and methanol (MeOH) at a concentration of 1.5 M were assessed. In the second experiment, five different Me_2_SO concentrations (1 M, 1.5 M, 2 M, 2.5 M and 3 M) and six different cooling rates (0.5 °C/min, 1 °C/min, 2.5 °C/min, 5 °C/min, 7.5 °C/min and 10 °C/min) were tested. Cryopreservation in this experiment was conducted in controlled-rate freezer. In the third experiment, the tissue size (50, 100 and 150 mg) as well as incubation time (15 or 30 min) in the cryoprotectants were tested. Lastly, the effects of sugar supplementation of cell viability was assessed by supplementing the cryomedium with either glucose, fructose, trehalose or sucrose at 0.1 or 0.3 M. Tissue pieces were 100 mg and an equilibration time of 30 min was used in this trial.

### Vitrification of testicular pieces

Vitrification was conducted by utilizing needle immersed vitrification (NIV) methodology as described by Marinović et al. (2018) [29]. In short, 50 mg testicular fragments were pinned to an acupuncture needle and immersed into two media prior to cryopreservation: the equilibration solution (ES) and the vitrification solution (VS). After 15 min immersion in ES and 1, 1.5 or 2 min immersion in VS (depending on the experiment), leftover medium was carefully adsorbed from the tissue by paper wipes and the needles were plunged into liquid nitrogen. After at least 1 day of storage, tissue pieces were warmed in three subsequent warming solutions at RT containing L-15 supplemented with 10% FBS and various concentrations of sucrose (WS1 – 3M; WS2–1 M; WS3 did not contain sucrose). Digestion and counting procedures were conducted as mentioned above.

Firstly, optimization of the vitrification protocol was conducted by testing the effects of three different equilibration solutions and three different vitrification solutions on spermatogonia viability similarly to brown trout oogonia [26]. Equilibration (ES1 – ES3) and vitrification (VS1 – VS3) solutions contained different combinations and concentrations of Me_2_SO, MeOH and PG (ES1: 1.5 M MeOH + 1.5 M PG; ES2: 1.5 M MeOH + 1.5 M M Me_2_SO ES3: 1.5 M PG + 1.5 M Me_2_SO; VS1: 1.5 M MeOH + 4.5 M PG; VS2: 1.5 M MeOH + 5.5 M Me_2_SO; VS3: 3 M PG + 3 M Me_2_SO). The extender used consisted of L-15 supplemented with 10% FBS, 25 mM HEPES and 0.5 M trehalose, while incubation time in VS was 1.5 min. In the second trial, the exposure of testicular fragments for 1, 1.5 or 2 min to two vitrification solutions (VS1: 1.5 M MeOH + 5.5 M Me_2_SO; VS2: 3 M PG + 3 M Me_2_SO) was tested. The ES consisted of 1.5 M PG + 1.5 M Me_2_SO while the extender composition was the same as in the previous trial.

## Spermatogonia transplantation

### Preparation of recipients for transplantaiton

Goldfish (*Carassius auratus*) spawners obtained from a local breeder were injected intraperitoneally with carp pituitary 24 h before collection of gametes at a dosage of 0.5 mg/kg for females and 1.5 mg/kg for males, and 12 h before collection of eggs at a dosage of 2.5 mg/kg for females only. Gametes were collected by abdominal massage and stored at 15 °C until fertilization (not more than 15 min). Eggs from five females were mixed together and fertilized with pooled milt from 10 males. Embryos were allowed to stick on a Petri dish and then transferred into an incubator at 23 °C. Embryos were injected under the blastodisc at the 2-cells stage without dechorionation with 100 mM solution of antisense *dead end* morpholino (*dnd-*MO) oligonucleotide according to Goto et al. (2012) (target sequence: CATCACAGGTGGACAGCGGCATGGA) using a M-152 micromanipulator (Narishige, Japan) and FemtoJet^®^ 4x microinjector (Eppendorf, Germany). Part of embryos injected with dnd-MO was co-injected with GFP-nos1 3‱UTR mRNA [31] to confirm successful depletion of primordial germ cells. Water was changed daily until hatching. Swim up embryos were fed with *Artemia* nauplii *ad libitum*.

### Transplantation

Transplantation was conducted into 11 dpf dnd-MO treated recipient larvae. Two different test groups were defined: (1) a recipient group in which fresh cells were injected and (2) a recipient group into which cryopreserved/thawed cells were injected. Due to the low vitrification effectiveness in common carp, only frozen cells were transplanted. In both cases, spermatogonia were enriched using 30% Percoll gradient according to Pšenička et al. (2015) [32]. Recipient larvae were anesthetized in a 0.05% Tricaine solution (SigmaAldrich) and approximately 5000 cells were injected into the peritoneal cavity of the recipients. Injected larvae (100 larvae per group) were transferred into fresh water and left to recover for two weeks and fed with *Artemia* nauplii. Germline chimeras were then transferred into aquaria and fed with an artificial food (Scarlet, Coppens). Water temperature was constantly held at 23±1 °C after the transplantation in order to prevent sex bias [33]. Control groups of intact control fish and morphants were exposed to the same rearing conditions as the experimental individuals were; however, no operations were conducted on them.

### Germline chimera identification

From each test group, 40 fish were euthanized by a tricaine overdose, decapitated and dissected 3 months post transplantation (BW: 5.6±2.3 g). Firstly, gonads were visually inspected for signs of gonadal development under a light microscope. Subsequently, gonads were excised and stored separately at –80 °C until RNA isolation. RNA was isolated using TRIzol reagent according to manufacturer instruction (Invitrogen). Isolated RNA was transcribed into cDNA using Transcriptor High Fidelity cDNA Synthesis Kit (Roche). Primers for RT-PCR were designed for goldfish and carp vasa gene and tested for specificity and to find suitable annealing temperature. Goldfish forward primer 5’- CGGCTAGCCTGAGAGATGAG, reverse primer 5’- GATCTCGATAACCCCGTTCA, expected amplicon size 196bp. Carp forward primer CGGCCGGCCGGAGAGATGAG, reverse primer GATCTGGATAACCCCATACA, expected amplicon size 200bp. Primers were diluted according to the manufacturer’s instruction. The reaction mixture for PCR contained 1 µl template cDNA, 0.5 µl forward and 0.5 µl reverse primer, 5 µl PPP Master Mix (Top-Bio) and 3 µl PCR H_2_O (Top-Bio). Reaction conditions were 35 cycles of 94 °C for 30 s, 58 °C for 30 s and 72 °C for 30 s. Products were analyzed on gel electrophoresis on 2% agarose gel on a UV illuminator.

### Statistical evaluation

All trials were tested in triplicates and for each replicate suitable control sample was used. Data is presented as mean ± standard deviation (SD). All percentage data was arcsine transformed prior to statistical analyses. One-way ANOVA with Tukey’s honest significant difference (HSD) test was applied in the trial with different cryoprotectants. Other trials were evaluated by two-way ANOVA with Tukey’s HSD. Significance level was set for all trial at *p* < 0.05. Statistical analysis was performed using STATISTICA v13.1 software (TIBCO Inc., Palo Alto, CA, USA).

## Results

### Freezing of carp testicular tissue

The highest viability in the first trial was observed using Me_2_SO (8.4%), since the use of other cryoprotectants resulted in significantly lower viability (Tukey’s HSD, p < 0.05; Fig. 1A). Combination of different Me_2_SO concentrations (1 to 3 M) and freezing rates (-0.5 to –10 °C/min) resulted in a wide range of viability among different combinations. Viability over 20% was recorded only when combining a –1 °C/min freezing rate with 2 M and 2.5 M Me_2_SO (Fig. 1B). Generally, slower cooling rates (-0.5 to –2.5 °C/min) resulted in higher viability in comparison to the faster cooling rates, while the resistance to the fastest cooling rate increased with higher Me_2_SO concentration. Additionally, the use of higher Me_2_SO concentrations and faster cooling rates resulted in higher amount of viable spermatozoa in cell suspensions indicating that optimal conditions for spermatozoa and spermatogonia are different.

**Fig. 1.**
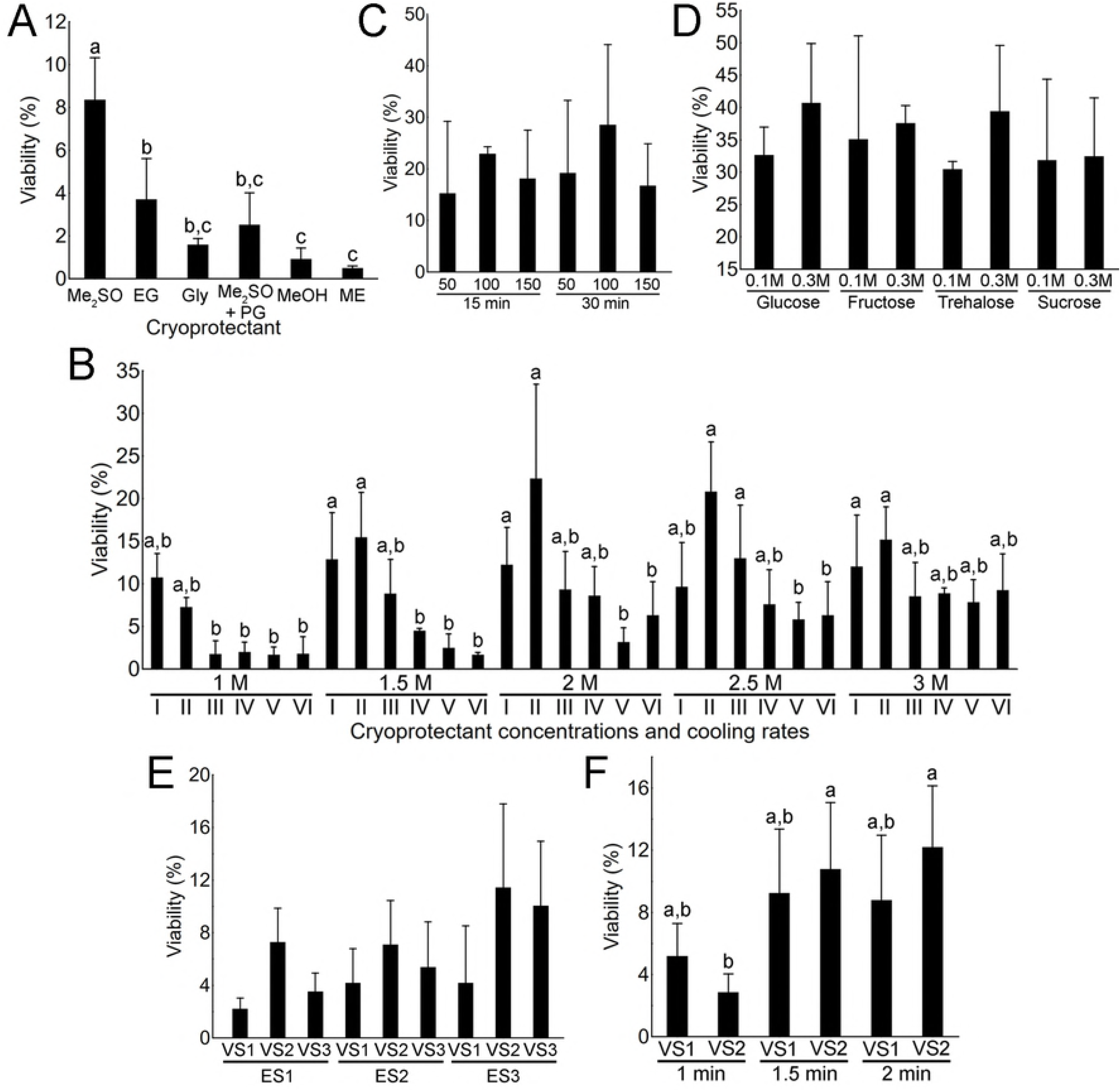
Optimization of the freezing (A-D) and vitrification (E, F) protocols for common carp spermatogonia. (A) Viability of spermatogonia after freezing with dimethyl sulfoxide (Me2SO), ethylene glycol (EG), glycerol (Gly), Me2SO and propylene glycol at ratio 1:1 (Me2SO+PG), methanol (MeOH) and metoxyethanol (ME). (B) The effects of Me2SO concentrations (1–3 M) and cooling rates of 0.5 (I), 1 (II), 2.5 (III), 5 (IV), 7.5 (V) and 10 (VI) °C/min on spermatogonia viability. (C) Viability of spermatogonia after exposing 50, 100 or 150 mg tissue fragments for 15 or 30 min to the cryomedium. (D) Effect of sugar supplementation of spermatogonia viability. Effects of different equilibration (ES) and vitrification (VS) solutions (E) and exposures (1–2 min) to different VS (F) on spermatogonia viability after NIV. All values are presented as mean ± SD. Different letters above the SD lines indicate statistical significance (Tukey’s HSD, p < 0.05), while the lack of such letters indicates the lack of statistical significance.

Exposure of tissue pieces of different sizes (50–150 mg) to cryoprotectants for variable periods of time (15 or 30 min) did not result in high variability. The highest viability was attained when equilibrating 100-mg tissue pieces for 30 min, however, statistical differences were not significant in comparison to other combinations (Tukey’s HSD, p > 0.05; Fig. 1C). Lastly, the supplementation of cryomedia with various sugars (glucose, fructose, trehalose and sucrose) in different concentrations (0.1 or 0.3 M) did not result in significant differences (Tukey’s HSD, *p* > 0.05; Fig. 1D). The highest viability of ∼ 40% was obtained when equilibrating 100 mg tissue pieces for 30 min in a cryomedia containing 2 M Me_2_SO, 0.3 M glucose, 1.5% BSA and 25 mM Hepes.

### Vitrification of carp testicular tissue

In the first vitrification trial, only the vitrification solutions displayed a significant effect on the viability of spermatogonia after warming (two-factor ANOVA, *p* < 0.05). Even though the average viability was higher when combining ES3 (containing 1.5 M PG and 1.5 M Me_2_SO) with either VS2 (containing 1.5M MeOH and 5.5 M Me_2_SO) or VS3 (containing 3 M PG and 3 M Me_2_SO), clear statistical differences could not be observed (Fig. 1E; Tukey’s HSD, *p* > 0.05). Therefore, VS2 and VS3 were used in the subsequent experiment. In the second trial, only the exposure times to the vitrification solutions had a significant effect on spermatogonia viability (two-factor ANOVA, p < 0.01). Only exposure for 1 min to VS2 (containing 3 M PG and 3 M Me_2_SO) yielded significantly lower viability rates compared to other groups (Fig. 1F; Tukey’s HSD, *p* < 0.05).

### Transplantation of cryopreserved spermatogonia

Due to the higher overall viability obtained by freezing (40.7 ± 9.2%) compared to vitrification (11.4 ± 4.9%), only spermatogonia frozen with the optimal protocol indicated above were transplanted alongside freshly isolated cells into the recipient goldfish larvae. As indicated above, recipient embryos were sterilized by injecting dnd-MO, and the success of sterilization was confirmed by fluorescent microscopy after co-injection with GFP-nos1 3‱UTR mRNA. All of the co-injected larvae displayed a successful depletion of recipient’s endogenous PGCs. *dnd*-MO injection affected the survival rates until the hatching stage compared to the untreated control during, however, survival after transplantation procedure and during ongrowing was comparable between assessed groups (Table 1).

**Table 1.** Recipient goldfish survival during the experiment.

| Treatment | No. of eggs fertilized | Survival 24 h (%) | Survival hatching (%) | Survival swim up (%) | Survival at transplantati on (%) | Survival 24 h pt | Survival 1 month pt | Survival 3 months pt |
| --- | --- | --- | --- | --- | --- | --- | --- | --- |
| Control | 235 | 199 (84.6) | 167 (71) | 151 (64.2) | 149<sup>a</sup> (63.4) | 149 | 143 | 142 |
| Morphants | 1060 | 719 (67.8) | 602 (56.8) | 554 (52.2) | 531* (50) | 100 (MO) | 92 | 92 |
|  |  |  |  |  |  | 96 (FC) | 88 | 88 |
|  |  |  |  |  |  | 93 (CC) | 87 | 86 |

pt – post transplantation
<sup>a</sup> no operation was conducted on the control group
\*MO treated goldfish larvae were divided for the transplantation into 3 groups, 100 larvae per group: MO – only *dnd*-MO treated fish, FC – fresh cells transplanted, CC – cryopreserved cells transplanted.

Success of transplantation was assessed three months after transplantation where the recipients were visually inspected for developing gonads after dissection, as well was by RT-PCR amplification of carp-specific *vasa* amplicons (Table 2, Fig. 2E). Firstly, during the visual inspection, none of the MO-treated control individuals showed any signs of developing gonads (Fig. 2B, 2B’) compared to the developing gonads observed in the non-treated controls (Fig. 2A, 2A’). The RT-PCR amplification of goldfish vasa amplicon additionally corroborated these findings as all assessed MO-treated controls and fish transplanted with cells were negative for goldfish *vasa* (data not shown) (Table 2).

**Fig. 2.**
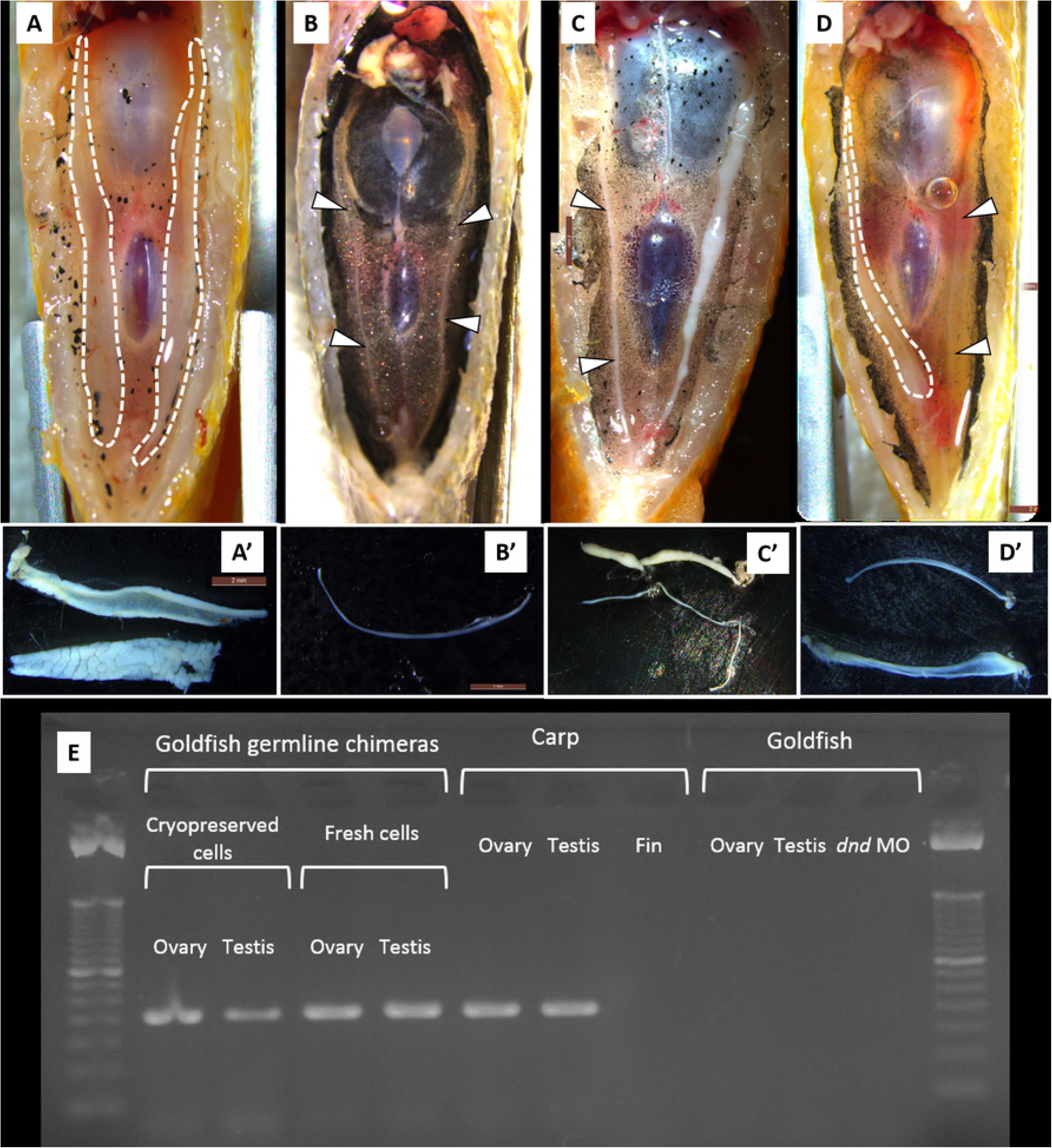
Detection of common carp spermatogonia incorporation and proliferation after interspecific transplantation into sterilized goldfish recipients. (A-D) Ventral view of dissected goldfish recipients. (A’-D’) Stereomicroscopic observation of the dissected gonads. (A, A’) Control fish displaying both gonads fully developed. (B, B’) dnd-MO treated goldfish displaying a lack of gonadal development. Development of testis (C, C’) and ovary (D, D’) after transplantation of common carp spermatogonia into dnd-MO sterilized goldfish recipients. Developed gonads are outlined with white dashed lines, undeveloped gonads are pointed out by white arrowheads. (E) RT-PCR amplification of common carp vasa amplicon in gonads of goldfish germline chimeras, as well as in control common carp gonads and fin and goldfish gonads.

**Table 2.** Summarized results of transplantation success and carp vasa RNA expression in germline chimeras evaluated 3 months post-transplantation.

| Treatment | Developed gonads/fish assessed | Testis/Ovary | Both gonads developed/one undeveloped | Carp *vasa* RT-PCR positive | Goldfish *vasa* RT-PCR positive |
| --- | --- | --- | --- | --- | --- |
| Control | 40/40 | 17/23 | 40/0 | 0 | 40 |
| Cryopreserved cells transplanted | 17/40 | 10/7 | 8/9 | 17 | 0 |
| Fresh cells transplanted | 21/40 | 14/7 | 9/12 | 21 | 0 |
| *dnd* MO treated | 0/40 | - | - | 0 | 0 |

Approximately 40% of the recipients injected with frozen/thawed carp spermatogonia displayed developing gonads (Table 2). Similarly, ∼ 50% of recipients injected with fresh spermatogonia displayed developing gonads. Developing gonads were either testes characterized by their while milky color (Fig. 2C, 2C’) or ovaries distinguishable by the presence of oocytes observable under higher magnification (Fig. 2D, 2D’) depending of the sex of the recipients; no intersex or individuals of indistinguishable sex were observed. Donor-derived origin of the germ cells within the developing recipient gonads was determined by RTPCR amplification of the carp *vasa* amplicon (Table 2; Fig. 2E). These results indicated that both fresh and frozen/thawed carp spermatogonia successfully migrated and incorporated into the goldfish gonads, as well as proliferated within the recipient gonads and produced later-stage germ cells of both sex.

## Discussion

In the present study, we have developed for the first time a cryopreservation methodology for common carp spermatogonia through freezing and vitrification of testicular tissue. The recovered germ cells were physiologically active since they were able to colonize and proliferate within the recipient gonads after inter-specific transplantation into *dnd*-MO sterilized goldfish recipients. Results of this study can serve as an alternative way for long-term preservation of common carp germplasm which can be recovered in a surrogate recipient through interspecific germ cell transplantation.

### Cryopreservation of testicular tissue

Me2SO was the most suitable cryoprotectant optimal for the freezing of common carp spermatogonia. Similar results were observed for salmonids [20,23,34] and cyprinids [25,35]. On the other hand, ethylene glycol which was optimal for freezing Siberian sturgeon (*Acipenser baerii*) spermatogonia [24] or propylene glycol suitable for freezing rainbow trout spermatogonia [20] were less effective in freezing of carp spermatogonia. Species-specific requirements and sensitivity to different cryoprotectants are therefore obvious and tailored optimized protocols are crucial. Cryoprotectant concentrations and cooling rates can be of crucial importance for freezing protocol optimization. These two parameters balance the rate of water efflux from the cell and its substitution for the cryoprotectant which will lower the freezing point and prevent the detrimental creation of intracellular ice [36]. Highest survival was recorded with 2 and 2.5 M Me_2_SO. Similar was reported in tench (*Tinca tinca*) and goldfish [25] where increased concentrations of cryoprotectants yielded higher spermatogonia viability. As for the cooling rates, lower cooling rates of –0.5 and –1 °C/min were more appropriate than the higher ones. Similar results were observed by Lee and Yoshizaki (2016) for the Manchurian trout (*Brachymystax lenok) [23]. The cause to this is most likely related to cell size: larger cells* generally require lower cooling rates and thus more time during cooling to enable water efflux out of the cell [37].

With regard to freezing of isolated cells or whole tissues, studies of Pšenička et al., 2016 and Marinović et al., 2016 [24,25] indicated slightly better results when freezing whole tissues. Cryopreservation of whole tissue is a more reasonable approach since gonadal tissue can be dissected, incubated in cryomedia, frozen to – 80 °C and then stored in liquid nitrogen within a timeframe of 2–3 hours (in case of slow cooling rate 1 °C/min). Cryopreservation procedure of isolated cells inherently takes longer since the tissue needs to be dissociated (and/or enriched for spermatogonia) which takes more time. Moreover, the use of this approach for germ cell transplantation could compromise its efficiency due to a high number of dead cells in the suspension and/or would optionally call for further purification of the suspension. However, when freezing whole tissues, attention needs to be paid to the size of the frozen tissue. In immature individuals or fish of small size, testicular tissue is generally small, therefore further fragmentation would not have any benefits [20,23,26]. On the other hand, when presented with large mature testes such as ones of common carp, its fragmentation is necessary. Trials of the present study did not display any effect of tissue size nor equilibration time on spermatogonia viability, however, we recommend that tissue pieces should be of reasonable size and should not surpass 100 mg (where an average of approximately 600,000 spermatogonia may be isolated from 100 mg of frozen/thawed testicular tissue).

During the recent decades, vitrification as a form of cryopreservation distinct to freezing where the formation of ice crystals is circumvented by using ultra-rapid cooling rates (up to 10^10^ °C/s) [38] has started to gain significant scientific attention. Several studies conducted fish, bird and mammalian testicular and ovarian tissue [26,29,39,40] indicated that vitrification offers advantages when compared to freezing with regard to cost- and time-effectiveness, low volumes of lN_2_ needed and other. Even though studies conducted on zebrafish (*Danio rerio*)[29], medaka (*Oryzias latipes*) [27] and honmoroko (*Gnathopogon caerulescens*) [28] testicular tissue display vitrification viability comparable to freezing viability, the present study demonstrated that viability of vitrified spermatogonia was approximately four-fold lower than the viability obtained through freezing. Similar was observed in goldfish and Wels catfish (*Silurus glanis*) (unpublished data). The main difference between the low and high vitrification viability species is the presence of spermatozoa. The proportion of spermatozoa to other testicular cells in mature common carp, goldfish and catfish testis is much higher than in zebrafish and medaka, while the vitrified honmoroko testicular tissue was immature and contained only early-stage germ cells. As vitrification of spermatozoa regularly displays lower quality compared to sperm freezing [41], high proportion of spermatozoa within the testicular tissue might form a selective barrier for the application of vitrification protocols in certain cases.

### Transplantation of cryopreserved tissue

In the present study, we demonstrated successful inter-specific transplantation of carp spermatogonia into goldfish recipients and the onset of the surrogate production technology between these two species. Inter-specific surrogate production offers several distinct advantages such as shortening of the reproduction time of long-term maturing species [17,32,42,43] or overcoming problems connected with poor reproduction performance [27]. The reasons for choosing goldfish in surrogate reproduction of carp are: (1) its small body size [44], (2) relatively fast maturation [45], (3) similar reproduction characteristics and management to carp [46], (4) short phylogenetic distance between carp and goldfish when even crossbreeds are viable [47], (5) available technology for recipient sterilization [30], and (6) proven resistance to diseases which represent a serious threat to carp such as Koi herpes virus [48].

Sterility of recipients is one of the main factors in successful application of surrogacy. Results of the MO-treatment in the present study correspond to previous reports of dnd-MO sterilization in goldfish [30], rainbow trout [49], sterlet [50] or zebrafish [51]. The lack fluorescent primordial germ cells after the co-injection of GFP-nos1 3′UTR mRNA and the absence of goldfish-specific *vasa* amplicons in recipient gonads in all tested individuals indicate that the sterilization was successful. Similarly, gene editing techniques using knock out approaches to target the *dnd* gene have successfully induced sterilization in Atlantic salmon (*Salmo salar)*[52]. However, Škugor et al., (2014) reported severe metabolic impairments in morphants, primarily in the sex hormone metabolism. Consequences of a lifetime absence of dnd induced by gene editing techniques need to be assessed, and different species might reach differently to such circumstances. For example, after intra-specific transplantation of zebrafish spermatogonia, only 5% of recipients sterilized through *dnd*-KO demonstrated donor cell incorporated [54]; on the other hand, incorporation rates were significantly higher when *dnd-* MO-KD recipients were used (unpublished results). Other sterilization techniques such as triploidization or hybridization usually applied in salmonids can be used for production of convenient recipients [17,34]. However, partial development of indigenous gonads can occur and alter production of donor derived gametes [43]. Therefore, sterility achieved through PGCs migration disruption via temporal RNA knockdown seems to be most convenient sterilization approach in case of goldfish [30,50,55], even when an immersion in vivo MO can be applied instead of microinjection [56].

Transplantation of both frozen/thawed and fresh spermatogonia into goldfish recipient larvae resulted in successful colonization and proliferation of carp germ cells within the recipient gonads. Incorporation rates were similar, thus demonstrating that frozen/thawed germ cells retain their physiological capabilities and can be used in surrogate production technology. Additionally, observed incorporation rates (40–50%) are within the range reported for various other species such as brown trout (*Salmo trutta*) (27%) and European grayling (*Thymallus thymallus*) (23–28%) germ cells transplanted into rainbow trout (*Oncorhynchus mykiss)* [57], allogenic transplantation in rainbow trout (60–70%) [49]; rainbow trout germ cells into masu salmon (*Oncorhynchus masou*) (68.5%) [22]. In many observed germline chimeras in our study, only one gonad was developed, while the other remained undeveloped, similarly to the *dnd-*MO treated controls. This phenomenon can be most likely attributed to the transplantation procedure, where germ cells are injected only from one side of the recipient’s body cavity. Therefore, cells can either stay near the place of injection and colonize only one gonad, or they can spread and migrate throughout the body cavity and colonize both gonads.

In the present study, both testes and ovaries were observed in the germline chimeras after transplantation of both cryopreserved and fresh spermatogonia. This offers the possibility for production of gametes of both sexes, and subsequently production of viable offspring originating even from a single donor. Sexual plasticity of germ cells after transplantation has been already described in several species when transplanted spermatogonia developed into both male and female gametes [20,43]. Sexual plasticity has a great importance when germ cells from extraordinary specimens are preserved. However, in goldfish, temperature can significantly affect sex differentiation and the final sex ratio. Thus, goldfish were constantly held at 23 ± 1 °C during the first month because temperature above 25 °C is known to trigger masculinization [33]. Observed sex ratio in goldfish chimeras was slightly biased in favour of testicular development. This can be attributed to the fact that transplanted germ cells will rather tend to respect their original sex. Anyway, temperature sensitivity gives a possible advantage to goldfish as a recipient, because sex can be modified very easily without hormonal treatment. However, further studies are necessary to elucidate biological pathways causing the switch from spermatogonia to oogonia and *vice versa* as well as the effects of the surrounding environment on exogenous cells. Future studies will focus on optimization of surrogate reproduction, reproductive characteristics of goldfish recipients as well cryopreservation of female germ cells which is crucial because it is currently the optimal way of preserving maternal genome.

## Conclusion

This study developed an optimal protocol for cryopreservation of carp male germ cells by freezing with subsequent restoration in goldfish as a surrogate host. Post-thaw viability of cryopreserved spermatogonia was improved over 40% through optimizing factors such as cryoprotectants, their concentrations, cooling rate, tissue size, incubation time and lastly sugars and their concentration. Importantly, our study showed that cryopreservation can be successfully performed without advanced cooling equipment when a commercially available cooling box placed in a –80 °C deep freezer can be used. Incorporation rates of fresh and cryopreserved spermatogonia were similar after inter-specific transplantation into surrogate goldfish and transplanted spermatogonia developed within both ovaries and testes. The donor derived origin was further confirmed by vasa gene expression in germline chimera gonads. The results could serve as an alternative strategy in breeding programs for male germplasm cryopreservation with subsequent recovery in goldfish hosts. Additionally, cryopreservation gives a possibility to synchronize and carry out transplantation according to the availability of hosts. The results of the present study can be used in combination with the hypothermic storage described by Lujić et al. (2018) where hypothermic storage is optimal for short-term storage of up to two weeks, while the freezing methodology developed in this study is optimal for long-term storage [58]. Further steps will be taken to develop a protocol for female germ cell cryopreservation as well to improve transplantation success using younger recipients or developing germ-less carp hosts for allogenic germ cell transplantation.

## Acknowledgements

We would like thank to staff at the Faculty of Fisheries and Protection of Waters as well staff at Department of Aquaculture at Szent István University.

